# Geographically structured genetic variation in the *Medicago lupulina* – *Ensifer* mutualism

**DOI:** 10.1101/117192

**Authors:** Tia L. Harrison, Corlett W. Wood, Katy D. Heath, John R. Stinchcombe

**Affiliations:** Department of Ecology and Evolutionary Biology, University of Toronto, 25 Willcocks St., Toronto, Ontario, M5S 3B2, Canada; Department of Plant Biology, University of Illinois, 505 S. Goodwin Avenue, Urbana, Illinois, 61801, United States; Centre for Genome Evolution and Function, University of Toronto, 25 Willcocks St., Toronto, Ontario, M5S 3B2, Canada

**Keywords:** mutualism, population genetics, genetic differentiation, gene flow, coevolution, invasion

## Abstract

Mutualisms are interspecific interactions affecting the ecology and evolution of species. Patterns of geographic variation in interacting species may play an important role in understanding how variation is maintained in mutualisms, particularly in introduced ranges. One agriculturally and ecologically important mutualism is the partnership between legume plants and rhizobia. Through characterizing and comparing the population genomic structure of the legume *Medicago lupulina* and two rhizobial species (*Ensifer medicae* and *E. meliloti),* we explored the spatial scale of population differentiation between interacting partners in their introduced range in North America. We found high proportions of *E. meliloti* in southeastern populations and high proportions of *E. medicae* in northwestern populations. *Medicago lupulina* and the *Ensifer* genus showed similar patterns of spatial genetic structure (isolation by distance). However, we detected no evidence of isolation by distance or population structure within either species of bacteria. Genome-wide nucleotide diversity within each of the two *Ensifer* species was low, suggesting limited introduction of strains, founder events, or severe bottlenecks. Our results suggest that there is potential for geographically structured coevolution between *M. lupulina* and the *Ensifer* genus, but not between *M. lupulina* and either *Ensifer* species.

## Introduction

The maintenance of variation within mutualistic interactions has been posed as a paradox because strong selection is expected to erode variation in mutualism related traits (Charlesworth, 1987, Heath and Stinchcombe 2013). One simple mechanism that could resolve this paradox is genetic differentiation between populations in mutualism traits, coupled with some gene flow between populations that introduces new variants. To evaluate this possibility, it is necessary to incorporate a geographic perspective into studies of mutualism to determine whether both interacting partners exhibit similar patterns of genetic structure on a landscape scale. Here we use whole genome sequencing and genotyping-by-sequencing to characterize patterns of genetic and geographic differentiation in the annual legume *Medicago lupulina* and its mutualistic rhizobial symbionts in their introduced North American range.

The potential for geographic structure to maintain variation in interspecific interactions is a core component of the geographic mosaic perspective on coevolution. A geographic mosaic describes a scenario where the structure and intensity of coevolution differs between populations, and is characterized by genetic differentiation between interacting populations at loci underlying coevolutionary traits, followed by gene flow that introduces new variants (Thompson, 2005). Adaptive genetic divergence in coevolutionary traits can arise from interactions with genetically differentiated populations of a single partner species or turnover of partner assemblages across a focal species' range (Nagano et al. 2014; Newman et al. 2015). Formal theory and meta-analyses suggest that gene flow between genetically differentiated populations can facilitate local adaptation in host-parasite systems by increasing within-population genetic variance (Gandon *et al.,* 1996; Greischar & Koskella, 2007; Hoeksema & Forde, 2008; Gandon & Nuismer, 2009). Although theoretical models indicate that geographic structure may similarly maintain genetic variance in mutualisms (Nuismer et al. 2000), empirical evidence in positive species interactions is scarce.

Gene flow between differentiated populations has the greatest potential to maintain variation in interspecific interactions when the scale of population differentiation in both partners is congruent. While there is strong evidence of geographic variation in mutualist quality (Thrall et al. 2000; 2007), and geographic covariation in traits mediating interactions (Anderson and Johnson 2007), we lack large-scale empirical examinations of population genetic structure in interacting mutualists. The few empirical studies that have examined parallel patterns of geographic structured genetic variation in both partners report conflicting results. Anderson et al. (2004), for example, found parallel patterns of isolation by distance between carnivorous *Roridula* plants and their hemipteran mutualists, albeit at different spatial scales, and suggested that these population genetic structures could facilitate co-adaptation within populations or regions. Parker and Spoerke (1998), in contrast, found no evidence of genetic structure in either the annual legume *Amphicarpea bracteata* or its nitrogen-fixing rhizobial symbionts. Béna et al. (2005) reported suggestive evidence of cospeciation between legumes in the genus *Medicago* and their rhizobial symbionts, but this genus-level analysis was not able to link phylogenetic patterns to coevolutionary processes that might have generated them.

In this study, we characterized and compared the geographic scale of genetic differentiation between the annual legume, *Medicago lupulina,* and its mutualistic nitrogen (N)-fixing bacteria, *Ensifer meliloti* and *E. medicae,* to determine whether gene flow between differentiated populations could maintain variation in this mutualism. Within the mutualism, legumes provide carbon (C)-based rewards and shelter for the bacteria (rhizobia), while bacteria fix atmospheric nitrogen (N) into plant-available forms. The *Medicago-Ensifer* mutualism is characterized by considerable coevolutionary genetic variation (Heath 2010, Heath et al. 2012), and several aspects of its biology suggest that there is substantial potential for geographic structure in both partners. *Medicago lupulina* is primarily a selfer, which reduces gene flow via pollen and promotes genetic differentiation. In addition, *M. lupulina* and *Ensifer* were introduced to North America relatively recently (approximately 300 years ago) and potentially multiple times (Turkington and Cavers 1979). Multiple and separate introductions of *M. lupulina* and *Ensifer* to North America could have created the necessary geographic structure to maintain mutualism variation in its introduced range.

One challenge in evaluating the potential for geographic structure to maintain genetic variation in mutualistic traits is that geographic structure might only be detected at specific genes involved the mutualism. Although genetic structure at genes involved in adaptation other aspects of the environment will contribute to population divergence, but these differences will not result in divergence in mutualism-related traits or genes, except in the case of linkage disequilibrium or pleiotropy. Therefore, a rigorous test of geographic structure in mutualisms would ideally quantify patterns of structure at symbiosis genes in addition to the whole genome. The mutualism between legumes and nitrogen-fixing rhizobia is especially promising in this regard. Genes mediating the interactions have been mapped (Wernegreen and Riley 1999; Barnett et al. 2001; Markmann and Parniske 2009; Reeve et al. 2010; Oldroyd 2013; Stanton-Geddes et al. 2013; Bravo et al. 2016; Klinger et al. 2016) and it is feasible to sequence entire bacterial genomes with next-generation sequencing rather than just a handful of markers. Using both whole genome sequences and sequences of symbiotic loci such as nitrogen fixation and nodulation genes previously shown to be involved in the symbiosis between *M. lupulina* and *Ensifer* (Wernegreen & Riley, 1999; Kimbrel *et al.,* 2013; Kawaharada *et al.,* 2015), we looked for signals of coevolution between legumes and their rhizobia genome wide and at individual symbiotic genes.

We asked three questions about the *M. lupulina* and *Ensifer* mutualism. First, is there geographic structure in the distribution of *E. meliloti* and *E. medicae* that could facilitate differentiation of *M. lupulina* populations? Second, do symbiotic genes in rhizobia indicate alternative patterns of coevolution compared to the whole genome? Finally, is population genetic structure in *M. lupulina* aligned with *Ensifer* genetic structure such that it could promote local- or regional-scale coevolution?

## Methods

### Study system

*Medicago lupulina* is a clover native to eastern Europe and western Asia and was introduced (potentially multiple times) to North America in the 1700s (Turkington and Cavers 1979). Today, *M. lupulina* is found across North America in temperate and subtropical areas, including all 50 states and most Canadian provinces (Turkington and Cavers 1979). It is primarily self-fertilizing and disperses seeds passively (Turkington and Cavers 1979; Yan et al. 2009) and consistent with this, previous studies in the native range (Europe and Asia) have found significant isolation by distance (Yan et al. 2009). *Medicago lupulina* is largely considered a weed, although it has been used as an inefficient fodder plant and was potentially introduced to North America along with agricultural crops.

Two species of *Ensifer,* free-living soil bacteria native to Europe and Asia, inhabit root nodules of *M. lupulina* : *E. meliloti* and *E. medicae.* Both can also associate with other *Medicago* species (Prévost and Bromfield 2003). It is assumed the *Ensifer* species arrived in North America with a *Medicago* species (Turkington and Cavers 1979). *Ensifer* species associate with plants at the start of a growing season, and at nodule senescence they dissociate from the plant, dispersing into soil, where they can be redistributed due to soil disturbance and water flow (i.e. no vertical transmission). Their genomes consist of a circular chromosome (3.65 Mb) and two plasmids (~1.3 Mb and ~1.6 Mb) (Galibert et al. 2001; Reeve et al. 2010). Recombination is restricted to *Ensifer* plasmids and horizontal gene transfer can occur between plasmids of different species of *Ensifer* (Bailly et al. 2006; Epstein et al. 2012; 2014). Many genes known to be involved in the mutualism, including *nif* and *nod* genes, are found on the plasmids of *E. meliloti* and *E. medicae* (Bailly et al. 2006; 2007), while housekeeping genes for general bacterial functions predominate the chromosome. Past studies have failed to detect significant genetic differentiation in *E. meliloti* and *E. medicae* populations in Mexico, suggesting high levels of gene flow in *Ensifer* populations (Silva et al. 2007).

### Field Sampling

We sampled *M. lupulina* individuals opportunistically from 39 populations across a wide geographic range in southern Ontario and the northeastern United States, a subset of *M. lupulina’s* introduced range (Supplemental Table 1). We randomly collected 2 to 10 plant individuals (spaced approximately 0.5 to 2 m apart) in late stages of their life cycle for both seeds and nodules. Seeds were collected in envelopes in the field and nodules were kept on the roots and placed in plastic bags at 4°C until processed. We obtained samples from 28 populations in southwestern Ontario (10 km to 300 km apart). To study large-scale geographic patterns, we sampled an additional 11 populations along a NW to SE transect from southern Ontario to Delaware, USA, separated by up to 820 km.

### Molecular Protocols

We extracted rhizobia samples from one field-collected nodule per plant, and used field-collected seeds to grow plant material for DNA extraction. Full details on plant growth conditions, bacterial plating and isolation procedures, and DNA extractions can be found in the Supplemental Materials (Appendix A). In brief, we isolated one bacterial strain per plant for whole genome sequencing using the MoBio UltraClean Microbial DNA Isolation Kit, and for the plants we isolated DNA from one individual per maternal line for genotyping-by-sequencing (GBS) according to the instructions of the Qiagen DNeasy Plant Tissue Mini Protocol. Genotyping-by-sequencing (GBS) is a high-throughput and cost-efficient method of sequencing large numbers of samples. GBS is similar to restriction site-associated sequencing (RAD-seq), and uses restriction enzymes to identify single nucleotide polymorphisms across the entire genome without sequencing the whole genome (Elshire et al. 2011). The GBS protocol is optimized for many different plant species, including *Medicago.*

We submitted 89 bacterial DNA samples to the Hospital for Sick Children (Toronto, ON, Canada) for library preparation and whole-genome sequencing on a HiSeq Illumina platform, using one lane and 2×100bp reads. For *Medicago,* we submitted 190 DNA samples to Cornell University (Ithaca, NY, United States) for GBS. The 190 DNA samples were distributed across two 96-well plates with 95 samples and one blank in each plate for the 96 multiplex GBS protocol. Cornell University prepared genomic libraries (Elshire et al. 2011) using a single digestion with EcoT22I (sequence ATGCAT). Samples were sequenced in two Illumina flow cells lanes.

### Bioinformatics and SNP discovery

We aligned forward and reverse rhizobia reads to the reference genome of *E. meliloti* strain 1021 (Galibert et al. 2001) (NCBI references chromosome AIL591688, plasmid a AE006469, plasmid b AL591985) and the *E. medicae* strain WSM419 (Reeve et al. 2010) (NCBI references chromosome 150026743 plasmid b 150030273, plasmid a 150031715, accessory plasmid 150032810) using BWA (Li and Durbin 2009) and Stampy (Lunter and Goodson 2011) with default parameters and the bamkeepgoodreads parameter. We assigned bacterial species using a combination of the percentage of reads mapping to one reference genome, and sequences at the 16S rDNA locus (NCBI gene references 1234653 and 5324158, respectively), which differs between *E. medicae* and *E. meliloti* (Rome et al. 1997). We used Integrative Genomics Viewer to visualize and check alignment quality (Robinson et al. 2011). In general, 69.99 – 94.02% (median 84.71%) of reads per sample mapped to the *E. meliloti* reference genome, and 69.32 – 92.48% (median 83.49%) mapped to the *E. medicae* genome.

In addition to creating a separate SNP file for each *Ensifer* species, we also created a single SNP file containing both *E. meliloti* and *E. medicae* (hereafter referred to as the “ *Ensifer* genus dataset”) to assess divergence between the two rhizobia. To create this file, we aligned all strains from both species to the *E. meliloti* reference genome and performed the same SNP discovery methods as performed on the *E. meliloti* species alignments (detailed below). We found shared polymorphisms between the two species and the two species were correctly identified in Structure (Supplemental Figure 1) and in Phylip (neighbour joining) (Supplemental Figure 2) using this data set (Pritchard et al., 2000, Felsenstein, 1989). To determine whether the reference genome we used influenced our results, we also aligned all the strains to the *E. medicae* reference genome. This analysis produced similar qualitative results (it correctly identified the two *Ensifer* species in Structure (Supplemental Figure 3)), so we used the *E. meliloti* alignments for the combined species SNP file for the rest of our analyses.

In *Ensifer,* we used PICARD tools to format, sort, and remove duplicates in sequence alignments. We applied GATK version 3 indel realignment and GATK Unified Genotyper SNP discovery on all bacteria alignments (McKenna et al. 2010) with ploidy set to haploid. We used the Select Variants parameter in GATK to select SNP variants only. We used standard hard filtering parameters and variant quality score recalibration on SNP discovery according to GATK Best Practices (DePristo et al. 2011; Van der Auwera et al. 2013). We filtered rhizobia SNPs for a minimum read depth (DP) of 20, a maximum DP of 226 for *E. meliloti* (230 for *E. medicae),* and a genotype quality (GP) of 30 using vcftools (Danecek et al. 2011). We removed indels and sites with more than 10% of missing data from both *E. meliloti* and *E. medicae* data files. We identified synonymous SNPs using SnpEff (Cingolani et al. 2012a) and SnpSift (Cingolani et al. 2012b), using reference files GCA_000017145.1.22 and GCA_000006965.1.22 (for *E. medicae* and *E. meliloti,* respectively) in the pre-built database. We used the ANN annotation parameter in SnpSift to identify SNPs as synonymous variants and missense variants.

We called *Medicago* SNPs in GBS samples by following the three-stage pipeline in the program Stacks (Catchen et al. 2011; 2013): cleaning raw data, building loci, and identifying SNPs. We trimmed reads to 64 bp and filtered reads by a phred score of 33, the default value for GSB reads sequenced on Illumina 2000/2500 machine. We built loci for *M. lupulina* using the *de novo* approach in Stacks (denovo_map command), setting the–m parameter at 5, the–M parameter at 1, and the-n parameter at 1. In the final stage of the pipeline, we identified SNPs under the populations command by setting the–m parameter at 5. We filtered SNPs by removing indels, removing sites with more than 10% of missing data, and removing sites that were less than 64 bps apart with vcftools (Danecek et al. 2011). We also excluded 9 SNPs with heterozygosity that was higher than expected under Hardy-Weinberg.

### *Analysis of* M. lupulina *and* Ensifer *genetic structure*

We tested whether genetic distance was correlated with geographic distance (isolation by distance) in *Medicago* and *Ensifer* using Mantel tests, implemented in R (R Core Team, 2016) with the ade4 package (Dray and Dufour 2007) using 100 000 randomizations. We estimated pairwise genetic distances between populations in *M. lupulina* and between individual samples in *Ensifer* because we sampled relatively few rhizobia from each population (1 – 3 samples). For *M. lupulina,* we used SNPs to calculate pairwise F_ST_ between populations in the program Genodive (Meirmans and van Tienderen 2004) using the population F_ST_ function and 1000 permutations, including only populations that had at least two individuals in F_ST_ estimates. We converted F_ST_ values to genetic distance values using F_ST_/(1-F_ST_) (Rousset 1997). In addition to calculating genetic distance between plant populations, we also used F-statistics to test for genetic differentiation between individuals hosting different species of bacteria, and to estimate population-level selfing rates [s = 2F_IS_/(1+F_IS_)] (Hartl and Clarke 1989). For *Ensifer,* we calculated Rousset’s genetic distance between strains in the program Genepop using the combined *E. medicae* and *E. meliloti* SNP dataset (Rousset 2008). To test for isolation by distance within *Ensifer* species, we repeated this procedure separately for *E. medicae* and *E. meliloti* data sets, and also computed separate tests of isolation by distance for the chromosome and plasmid to assess structure at different components of the *Ensifer* genome.

Second, we tested for spatial genetic autocorrelation of allele frequencies in *M. lupulina,* in the *Ensifer* genus, and separately in each *Ensifer* species using GenAlEx v.6.5 (Peakall & Smouse, 2006, 2012). This analysis tests against the null hypothesis that genotypes are randomly distributed in space. We binned individuals into 8 distance classes of 100km for the *M. lupulina* and *Ensifer* genus analyses, and into 4 distance classes of 200km for the separate analyses of each *Ensifer* species, because our sample sizes were smaller for the latter two analyses. We tested for significant spatial autocorrelation by permuting individuals among geographic locations (N_permutations_ = 999) and placed confidence limits on our estimates of spatial autocorrelation using 1000 bootstrap replicates.

Finally, we tested for a geographic pattern in the distribution of the two *Ensifer* species. Because our sampling transect ran from northwest to southeast, we created a single variable representing increasing longitude and decreasing latitude by extracting the first principal component (“PC1”) from the latitude and longitude coordinates of our collection sites. The PC1 axis captured 90.79% of the variance in geographic location among our collection sites. We regressed the proportion of *E. meliloti* samples in a site on PC1 to identify the relationship between *Ensifer* species proportion and geographic location. (R Core Team, 2016). To assess whether spatial autocorrelation of plant samples impacted the results of this analysis, we randomly removed 17 Ontario populations and re-ran our analysis on the remaining 11 Ontario populations and the 11 American populations. We repeated this procedure 100 times, and obtained qualitatively similar results to the full dataset in all cases (P ≤ 0.0001 in all cases), indicating that the geographic pattern in the distribution of the bacteria species is robust to our uneven geographic sampling.

### Analysis of rhizobial nucleotide diversity and symbiosis genes

We next looked for genetic variation between strains within the same *Ensifer* species. Specifically, we assessed nucleotide diversity within *Ensifer* species by calculating the average pairwise nucleotide differences (π) between rhizobial samples. We extracted average pairwise nucleotide differences from *Ensifer* vcf files using a custom Python script (Python Software Foundation, 2010). We averaged all pairwise nucleotide differences across strains to obtain π, and divided it by the number of loci (variant and non-variant) called by GATK to obtain per site values. We calculated π for the range-wide sample, and repeated this calculation including only individuals collected from southern Ontario, which are in close proximity and more likely to experience similar environmental (and potentially selective) conditions. We calculated π separately for the *Ensifer* chromosome and two plasmids and for synonymous and nonsynonymous SNPs in both species of *Ensifer.*

In addition to calculating nucleotide diversity at the genome-wide scale, we also calculated nucleotide diversity for individual genes known to be involved in the symbiosis between *M. lupulina* and *Ensifer* species (Wernegreen and Riley 1999): nodulation genes *nodA, nodB,* and *nodC*; and nitrogen fixation genes *nifA, nifB, nifD, nifE, nifH, nifK, nifN,* and *nifX* (NCBI gene reference numbers given in Supplemental Table 2). Previous research has also identified pathogen type III effector genes as important genes in host infection (Kimbrel et al. 2013), so we calculated nucleotide diversity for two type III effector loci in *E. medicae* (Reeve et al. 2010). In addition, there is evidence that bacterial exopolysaccharides are involved in nodule formation and rhizobia infection (Kawaharada et al. 2015). We estimated nucleotide diversity in one gene (exoU glucosyltransferase) that produces exopolysaccharides in *E. meliloti* (Finan et al. 2001).

To further characterize diversity among rhizobia samples and more specifically assess how rare polymorphisms are in the rhizobia samples, we also constructed minor allele frequency spectra of the *E. medicae* of *E. meliloti* data. We removed 100% of missing data from the *E. medicae* and *E. meliloti* vcf files before calculating allele frequencies for synonymous and non-synonymous SNPs using vcftools (Danecek et al. 2011). We extracted the least frequent alleles from the *Ensifer* vcf files and constructed histograms of *E. medicae* and *E. meliloti* minor allele frequencies in R using the plotrix package.

### Comparison of M. lupulina and Ensifer genetic structure

To determine whether *M. lupulina* and *Ensifer* exhibited similar patterns of isolation by distance, we tested whether pairwise genetic distances between *M. lupulina* individuals were correlated with pairwise genetic distances between their rhizobia, using a Mantel test with 100 000 randomizations. We used *Ensifer* genus dataset (combined *E. meliloti* and *E. medicae*) to estimate individual genetic distance in *Ensifer.*

We estimated population structure among samples in *M. lupulina* and in the *Ensifer* genus using a combination of InStruct (Gao et al. 2007) and Structure (Pritchard et al. 2000). For *M. lupulina,* we tested for a maximum population value (K) of 5 under the admixture and population selfing rate model (v = 2) in the program InStruct (which allows for population assignments in selfing organisms). We ran 2 chains for each K value with 500 000 000 repetitions and a burnin of 200 000 000 and included no prior information. All other InStruct parameters were kept at default values. The Gelman-Rudin statistic confirmed that convergence among chains was achieved. We used the Deviance Information Criteria (DIC) to select the value of K that provides the best fit to the data. We post-processed Structure runs using CLUMPP (Jakobsson and Rosenberg 2007) and made plots using Distruct (Rosenberg 2004).

Before we estimated population structure in rhizobia strains using Structure, we first estimated recombination among the samples. The Structure model assumes that loci are not in linkage disequilibrium within populations (Pritchard et al. 2000), which is likely to be untrue for non-recombining regions like the *Ensifer* chromosome (Bailly et al. 2006). We used the program ClonalFrame (Didelot and Falush 2007) to estimate *ρ/θ* (number of recombination events/number of mutation events). We used VCFx software (Castelli et al. 2015) to convert our *Ensifer* genus vcf file of combined *E. meliloti* and *E. medicae* SNPs to an aligned fasta file – the input format for ClonalFrame. We performed 2 runs of ClonalFrame with 100 000 iterations and removed 50 000 as the burnin. We checked for convergence using Gelman and Rubin’s statistic. ClonalFrame identified a sufficiently high rate of recombination (*ρ/θ* = 1.0021) among *Ensifer* samples to justify Structure analysis. In Structure (Pritchard et al. 2000), we performed 5 runs with 200 000 iterations and discarded 100 000 for the burnin. We tested for a maximum K of 5 under a model of admixture and correlated allele frequencies. We used StrAuto to automate Structure processing of samples (Chhatre 2012). All summary statistics (alpha, F_ST_, and likelihood) stabilized before the end of the burnin. We then used Structure Harvester to detect the inferred K in the likelihood data generated by the Structure tests (Earl 2012), using the deltaK approach (Evanno et al. 2005). Structure runs were post-processed and plotted as described above.

To assess phylogenetic congruence between *Medicago* and *Ensifer,* we estimated phylogenetic relationships among individuals for the plant and the rhizobia by constructing maximum likelihood trees in RAxML (Stamatakis 2014). We used the GTRGAMMA function with 100 bootstraps to build our trees. Because we used SNP alignment files without invariable sites included we used the ASC_ string to apply an ascertainment bias correction to our data set. We built a maximum likelihood tree for *M. lupulina* samples and the *Ensifer* genus (based on the combined *E. medicae* and *E. meliloti* SNP data). We then used the cophyloplot function and the dist.topo function in phangorn (Schliep 2011) in R to visualize the two trees and calculate topological distance between the trees. We also estimated separate neighbour joining trees for the *Ensifer* chromosome and two plasmids using the ape package (Paradis et al. 2004) in R to compare structure at different components of the *Ensifer* genome.

## Results

### Medicago lupulina *genetic structure*

The *M. lupulina* sample of 190 individuals comprised 39 populations and 2349 SNPs, and exhibited a significant signal of isolation by distance (Figure 1). The positive relationship between geographic distance and genetic distance indicates that populations farther apart are more genetically different than populations located close together. Population-level selfing rates (Supplemental Table 3) were quite high on average (s = 0.813), which may contribute to isolation by distance in *M. lupulina.* F_ST_ between *M. lupulina* individuals hosting *E. medicae* and individuals hosting *E. meliloti* was low (0.0190 ± 0.0001) but significant (p = 0.0010).

**Figure 1.**
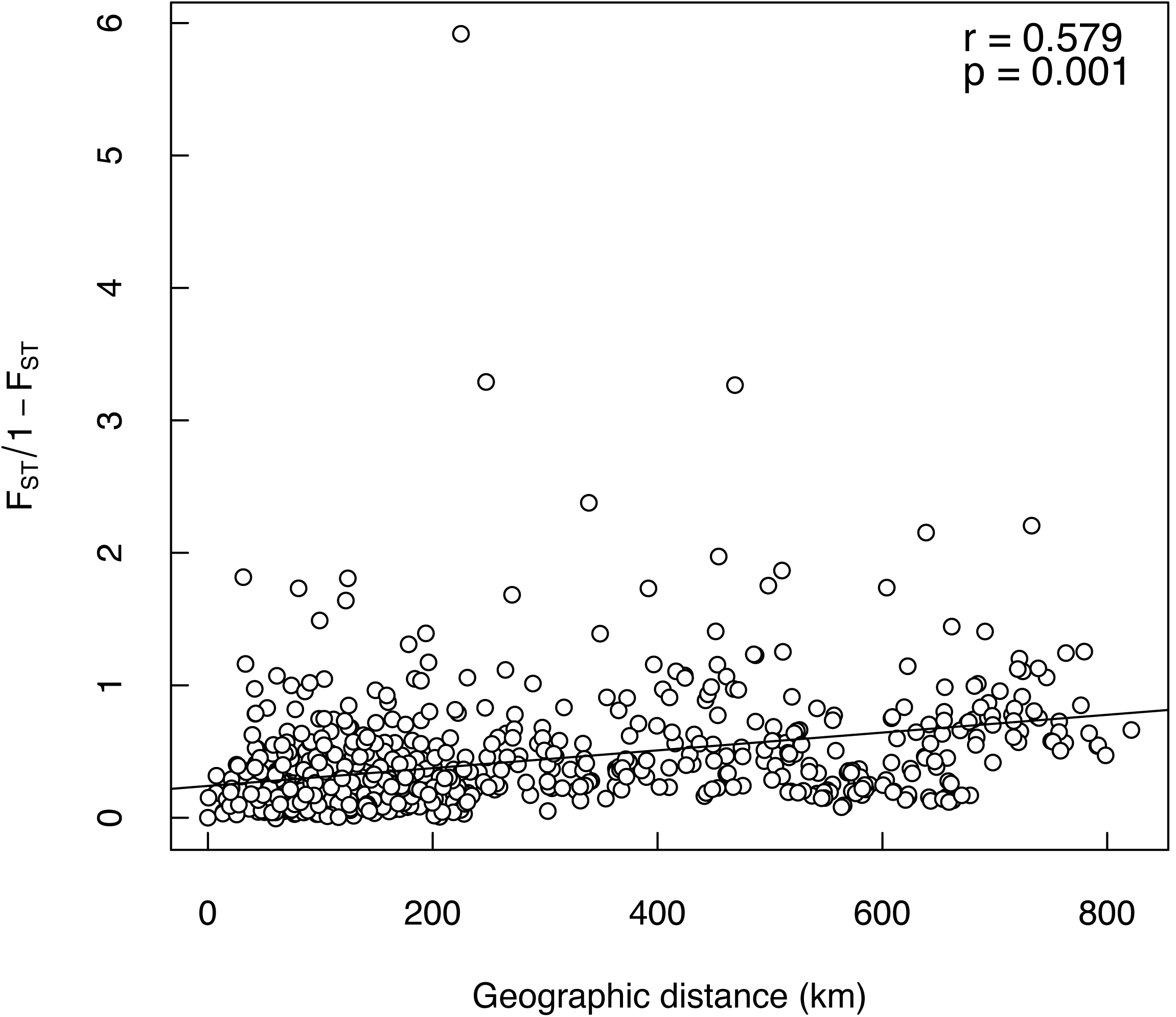
Relationship between geographic distance and genetic distance in *Medicago lupulina* populations. Each point represents a pairwise population comparison. One population was removed from the *M. lupulina* data set because it produced an abnormally high genetic distance value when compared pairwise with other populations (population 11).

There was significant spatial autocorrelation of allele frequencies in *M. lupulina* (Supplemental Table 4, Supplemental Figure 4A). We found a positive spatial autocorrelation between individuals located within approximately 200km of each other (r ≥ 0.04, P = 0.001), indicating that geographically proximate individuals are more closely related than the null expectation. We found a negative spatial autocorrelation between individuals located farther than 300km from each other (r ≤ −0.01, P = 0.001), indicating that geographically distant individuals are less closely related than the null expectation. These results are consistent with the pattern of isolation-by-distance reported above.

### Ensifer *genetic structure*

We assigned 50 rhizobia samples to *E. meliloti* and 39 samples to *E. medicae;* summary statistics on sequencing can be found in Supplemental Tables 5 and 6. The 39 *E. medicae* samples were distributed among 24 populations. In this dataset, we discovered 1081 SNPs, of which 678 were synonymous and 209 non-synonymous. The 50 *E. meliloti* sample were distributed among 28 populations, but contained approximately half the number of SNPs that *E. medicae* did (554: 234 synonymous and 176 non-synonymous). Our *Ensifer* genus dataset (combining both *E. meliloti* and *E. medicae*) contained a total of 89 samples and 476 SNPs; this dataset contained fewer SNPs than either the *E. medicae* or *E. meliloti* datasets because it only includes sites that were genotyped in both species.

Population composition of bacteria species changed significantly with longitude and latitude. When we regressed the proportion of plants associated with *E. meliloti* on PC1, which represented increasing longitude and decreasing latitude of our sampling locations, we found a positive significant relationship (F_1,37_ = 15.804, P < 0.001). Populations in the southeast contained higher proportions of *E. meliloti* while populations in the northwest contained higher proportions of *E. medicae* (Figure 2).

**Figure 2.**
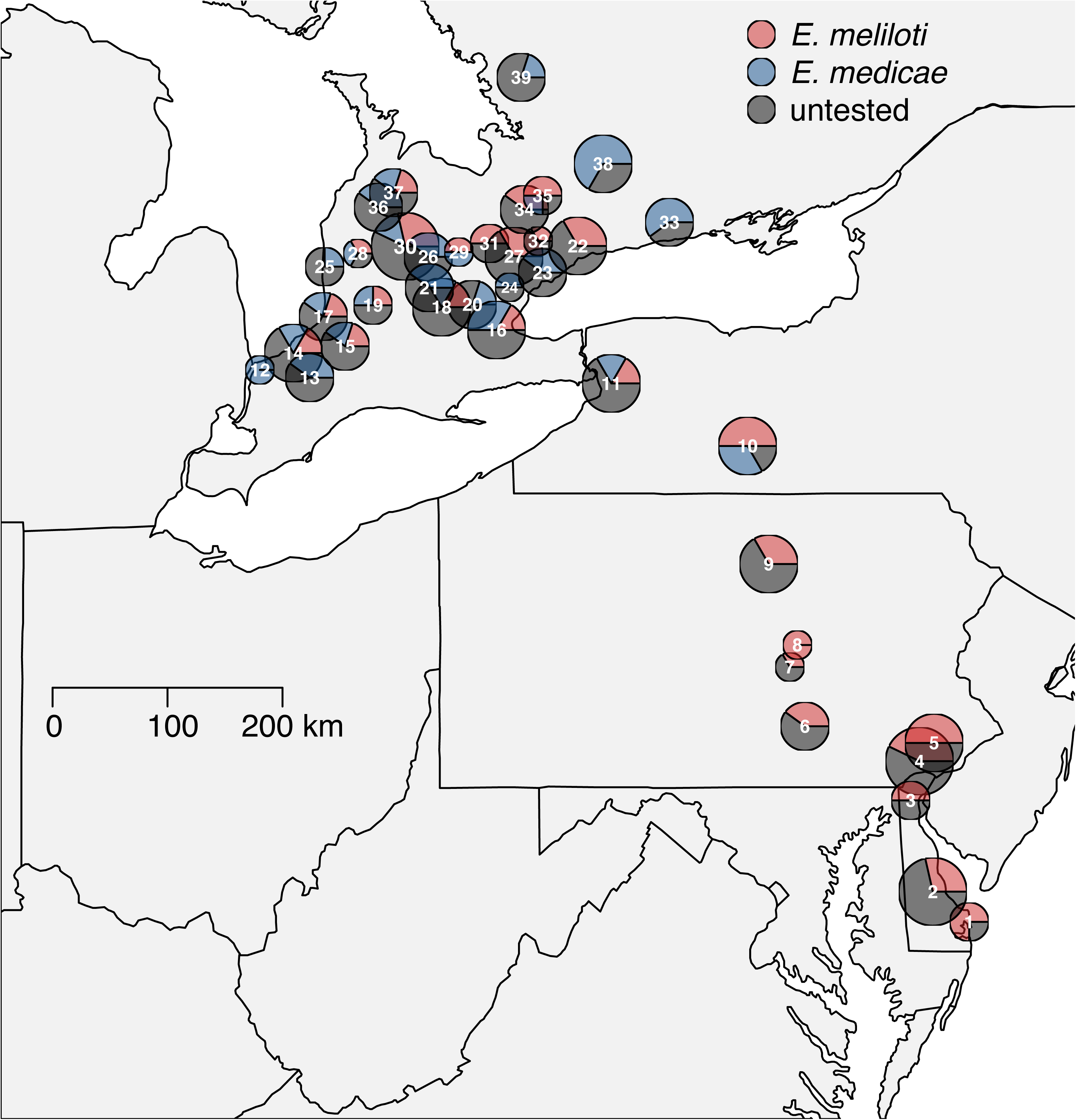
Population composition of *E. meliloti* and *E. medicae* in *M. lupulina* populations in North America. Radius of circle corresponds to the number of *M. lupulina* samples collected in the population. Pie charts represent the proportion of plants from each population that were hosting *E. meliloti* (red), *E. medicae* (blue), and an unidentified rhizobia species (grey). Populations are numbered from south to north.

We found a significant signal of isolation by distance in our *Ensifer* genus data set (Figure 3a), as expected given the geographic cline in their frequencies (Figure 2). We failed to detect isolation by distance within either *Ensifer* species using whole-genome data (Figure 3b and c). There was also no significant isolation by distance when we performed this analysis using only SNPs from the bacterial chromosome and plasmids in either *Ensifer* species (*E. medicae:* 0.23 < p < 0.65; *E. meliloti:* 0.9 < p < 0.96).

**Figure 3.**
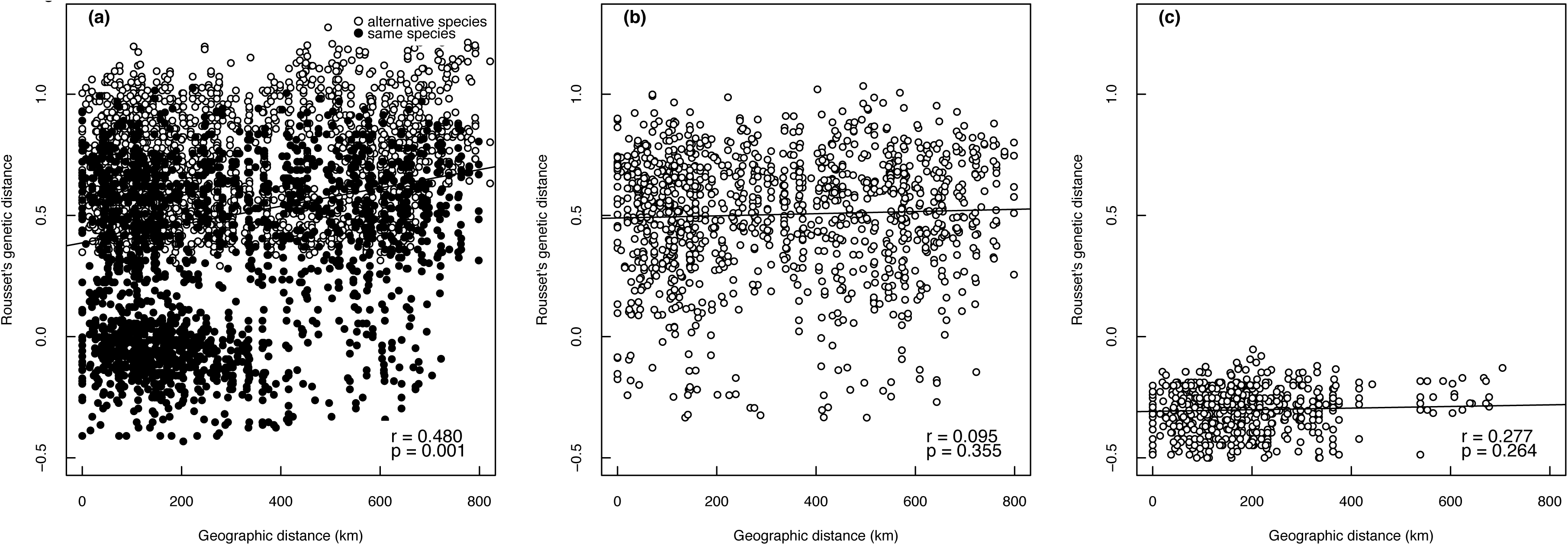
Relationship between geographic distance and Rousset’s individual genetic distance in a) total *Ensifer* genus data set (*E. meliloti* and *E. medicae*), b) *E. meliloti,* and c) *E. medicae.* Each point represents a pairwise individual comparison.

There was significant spatial autocorrelation in allele frequencies in the *Ensifer* genus (Supplemental Table 4, Supplemental Figure 4B). We found a positive spatial autocorrelation between individuals located within approximately 300km of each other (r ≥ 0.02, P ≤ 0.015), and a negative spatial autocorrelation between individuals located at least 600km from each other (r ≤ −0.05, P ≤ 0.004). These results are consistent with the pattern of isolation-by-distance reported above, in which geographically proximate individuals are more genetically similar (in this case, of the same species) and geographically distant individuals are more genetically dissimilar (i.e., of alternate bacterial species) than expected by chance. By contrast, there was no significant spatial autocorrelation of allele frequencies within either *Ensifer* species when the two species were analyzed separately (Supplemental Table 4, Supplemental Figures 4C and 4D).

### Ensifer *nucleotide diversity and symbiosis genes*

Genome wide nucleotide diversity values were extremely low within both *Ensifer* species in our full range data set and reduced data set in Ontario (Table 1). Symbiosis genes were particularly conserved (Table 2). We discovered only one to two SNPs in the *nodC* nodulation gene in both species of *Ensifer. NodA* and *nodB* genes contained no SNPs in either *E. medicae* or *E. meliloti.* In addition, *nifH* was the only nitrogen fixation gene that contained SNPs in both *E. medicae* and *E. meliloti*; *nifE* in *E. medicae* was the only other nitrogen fixation gene with a nucleotide diversity value greater than zero. We detected no SNPs in *E. medicae* type III effector genes or exopolysaccharide genes in *E. meliloti,* which are known to be involved in nodule formation and rhizobia infection (Kawaharada et al. 2015).

**Table 1.** Nucleotide diversity (mean π) of *E. medicae* and *E. meliloti* for different structures of the *Ensifer* genome.

| | $\pi$ synonymous | $\pi$ non-synonymous |
| --- | --- | --- |
| **Full range sample** |  |  |
| *E. medicae* |  |  |
| Chromosome | 0.0006108 | 0.0001117 |
| pSMED01 | 0.0010950 | 0.0002371 |
| pSMED02 | 0.0025284 | 0.0010754 |
| *E. meliloti* |  |  |
| Chromosome | 0.0001349 | 0.0000312 |
| pSymA | 0.0005108 | 0.0003873 |
| pSymB | 0.0001449 | 0.0000362 |
| **Southern Ontario sample** |  |  |
| *E. medicae* |  |  |
| Chromosome | 0.0004844 | 0.0000931 |
| pSMED01 | 0.0021592 | 0.0008977 |
| pSMED02 | 0.0009104 | 0.0002091 |
| *E. meliloti* |  |  |
| Chromosome | 0.0001324 | 0.0000283 |
| pSymA | 0.0005056 | 0.0003586 |
| pSymB | 0.0001338 | 0.0000383 |

**Table 2.** Nucleotide diversity (mean π) on nodulation genes and nitrogen fixation genes located on *E. medicae* and *E. meliloti* plasmids.

| | Number of SNPs | $\pi$ |
| --- | --- | --- |
| *E. medicae* |  |  |
| nodA | 0 | 0 |
| nodB | 0 | 0 |
| nodC | 2 | 0.0000761 |
| nifA | 0 | 0 |
| nifB | 0 | 0 |
| nifD | 0 | 0 |
| nifE | 1 | 0.0000359 |
| nifH | 3 | 0.0004896 |
| nifK | 0 | 0 |
| nifN | 0 | 0 |
| nifX | 0 | 0 |
| type III effector 4319 | 0 | 0 |
| type III effector 1279 | 0 | 0 |
| *E. meliloti* |  |  |
| nodA | 0 | 0 |
| nodB | 0 | 0 |
| nodC | 1 | 0.0002755 |
| nifA | 0 | 0 |
| nifB | 0 | 0 |
| nifD | 0 | 0 |
| nifE | 0 | 0 |
| nifH | 1 | 0.0001262 |
| nifK | 0 | 0 |
| nifN | 0 | 0 |
| nifX | 0 | 0 |
| exoU glucosyltransferase | 0 | 0 |

Minor allele frequency spectra showed that most minor alleles were very low in frequency in *E. meliloti* and *E. medicae* (Supplemental Figure 5). Minor alleles are all quite rare in *E. medicae* as almost all the alleles were below 5% in frequency. Minor allele frequencies in *E. meliloti* had more variation across the different frequency bins compared to *E. medicae* but still most of the alleles were low in frequency (5%).

### *Comparison of* M. lupulina *and* Ensifer *genetic structure*

We found a significant positive relationship between *M. lupulina* genetic distance and *Ensifer* genetic distance (Figure 4). The positive relationship indicates that as genetic divergence between plant populations increased, so did genetic differentiation between their associated rhizobia.

**Figure 4.**
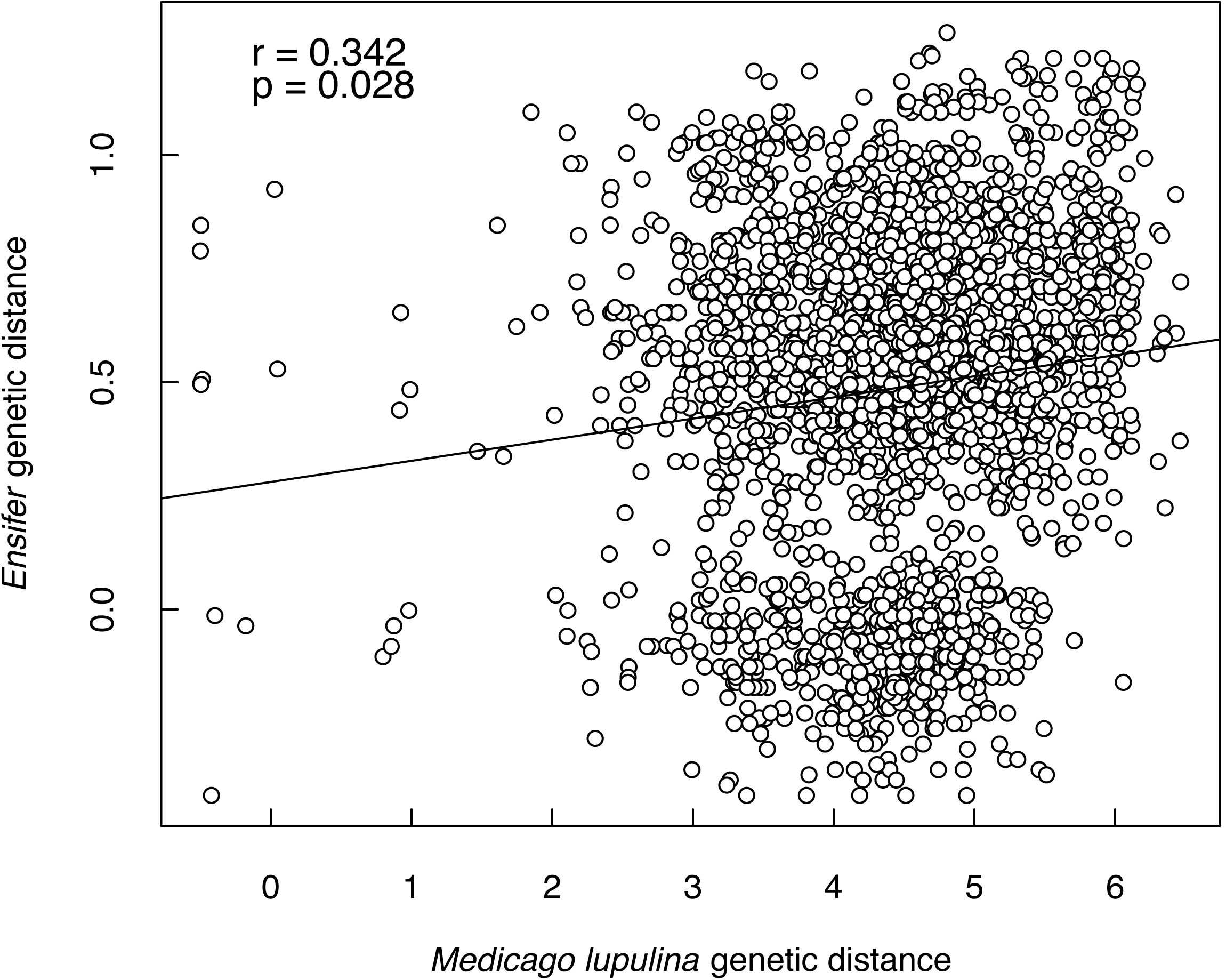
Relationship between *M. lupulina* individual genetic distance and *Ensifer* individual genetic distance. Each point represents a pairwise comparison between the genetic distance between two *M. lupulina* individuals and the genetic distance between their two corresponding rhizobia strains.

We compared population assignments in *Ensifer* samples to population assignments in their specific *M. lupulina* individual hosts. We identified two genetic clusters within *M. lupulina* using Instruct (Figure 5a), using the Deviance Information Criteria (DIC) to determine which value of K provided the best fit to the data. There is a weak geographic trend of northern *M. lupulina* individuals associated with the purple cluster, and southern *M. lupulina* individuals associated with the yellow cluster. Similarly, Structure Harvester identified 2 clusters within the *Ensifer* genus data set, corresponding to *E. medicae* and *E. meliloti* (Figure 5b). All *E. meliloti* samples were assigned to the red population and all *E. medicae* samples were assigned to the blue population.

**Figure 5.**
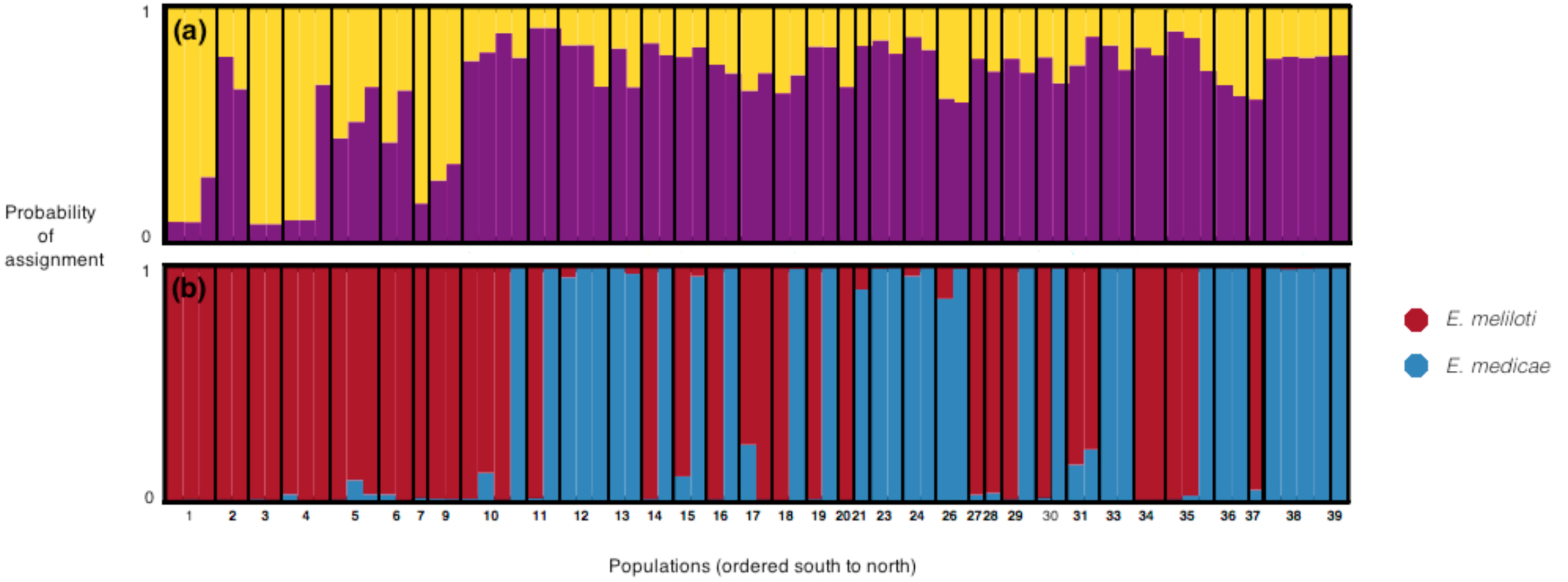
Population structure of a) *M. lupulina* and b) *Ensifer* genus. Black lines represent population divisions in the sample. Geographic population numbers are listed on the x-axis and are ordered from south populations to north populations.

The maximum likelihood trees of *M. lupulina* and *Ensifer* show extensive mismatching between tree tips (Figure 6). Plants hosting *E. medicae* and plants hosting *E. meliloti* did not group together on the *M. lupulina* tree. In addition, topological distance (the number of partitions that differ between the two trees) between the two trees was high (topological distance = 140, total partitions = 140, percent differences in bipartitions between trees = 100 %). It is important to note that both trees had low bootstrap support at internal nodes. The *Ensifer* tree had particularly low bootstrap at nodes within *Ensifer* species (which could be a result of the low genetic diversity within *Ensifer* species). Therefore, mismatches between *M. lupulina* and *Ensifer* at the tree tips is likely due in part to error associated with clade assignments.

**Figure 6.**
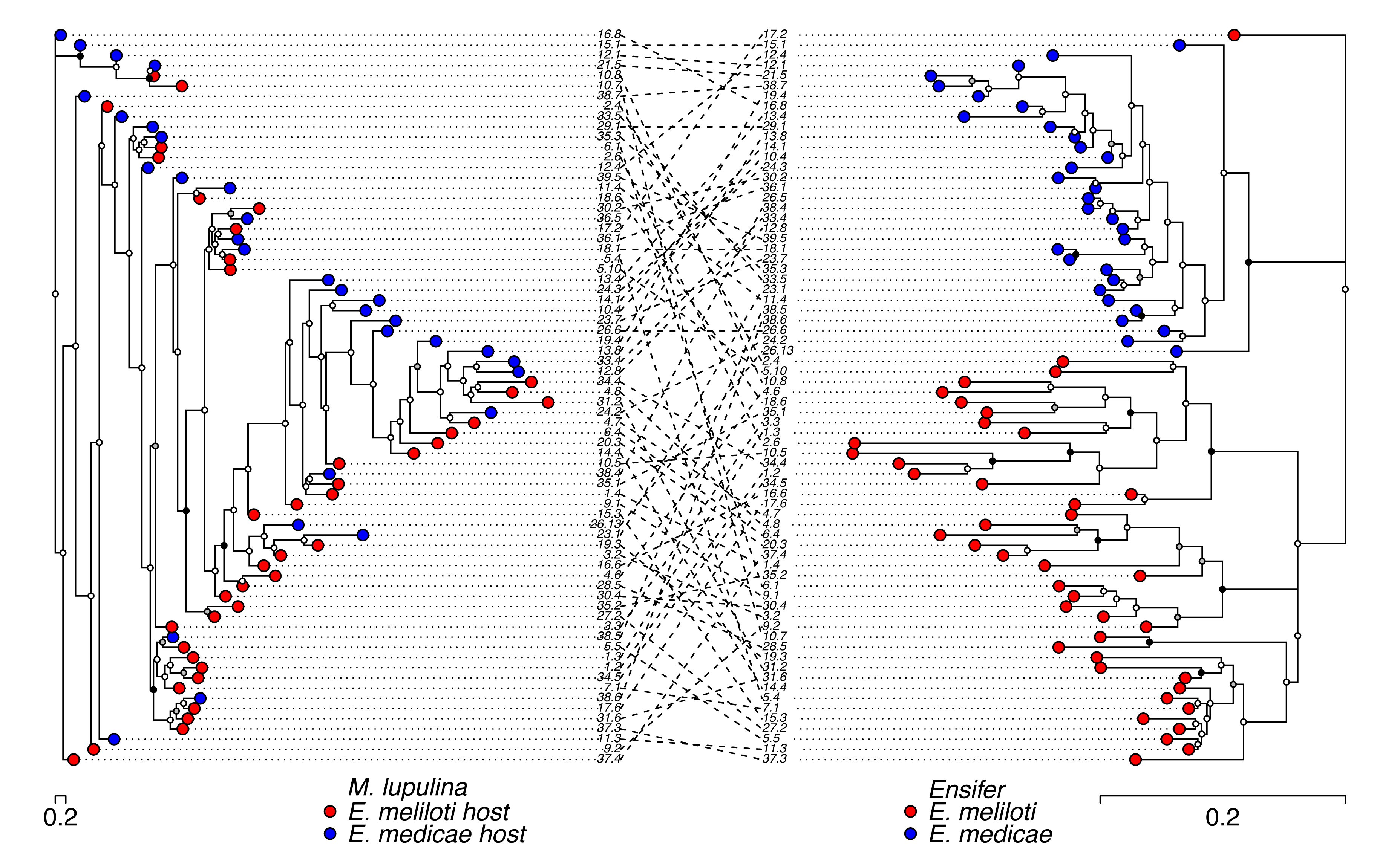
Phylogenetic analysis of *M. lupulina* (left) and *Ensifer* (right) estimated using genome wide SNPs. Maximum likelihood trees with posterior support given at each node. Circles at nodes indicate varying bootstrap support with the colours white (< 75%), grey (>75 < 90%), and black (>90%). Scale bar represents the genetic distance between individuals. Number codes represent populations and individuals within populations. Individuals are also labeled for which rhizobia species they were associated with in the sample (left tree) or which rhizobia species (right tree) they were identified as (red = *E. meliloti* and blue = *E. medicae*).

Groupings in the maximum likelihood tree of *M. lupulina* samples did not necessarily corresponded to groupings of geographic populations. The tree topology also showed large genetic distance between individuals. The tree topology for the *Ensifer* genus showed *E. medicae* and *E. meliloti* clearly separated into two groups (Figure 6). Groupings of *Ensifer* samples in the tree did not necessarily associate with geographic location, even when we constructed separate trees for the *Ensifer* chromosome and two plasmids. The chromosome and plasmid trees differed appreciably (Supplemental Figures 5 and 6). In general, the *Ensifer* tree had lower genetic distance between individuals when compared to the *M. lupulina* tree.

## Discussion

Our primary goal was to characterize and compare the spatial scale of genetic differentiation in the *M. lupulina* and *Ensifer* mutualism in a portion of its introduced range in eastern North America. The dominant picture that emerges from these analyses is that there is geographic structure in the *Ensifer* genus but very little genetic variation within each *Ensifer* species.

Therefore, the geographically structure of genetic variation, and potential for coevolution in this mutualism, appears mainly to be due to *M. lupulina* interacting with different bacterial species across its range, rather than genetically variable strains within a single bacterial species. Three major results emerged from our analyses, which we discuss in turn below: (1) The geographic turnover of *Ensifer* species composition in eastern North America, (2) The overall paucity of genetic variation within both *Ensifer* species, despite an extensive collection across a wide geographic range, and (3) Somewhat concordant geographic patterns of genetic variation in *M. lupulina* and the *Ensifer* genus.

### *Geographic turnover of* Ensifer *assemblages and low genetic variation within* Ensifer *species*

We showed that there is strong geographic structure in *Ensifer* mutualism assemblages in eastern North America. The rhizobia species *E. medicae* is more common in southern Ontario, with *E. meliloti* more common in northeastern and mid-Atlantic regions in the United States. Our results corroborate previous work, which found that *E. medicae* is more abundant in southern Ontario than other *Ensifer* species (Prévost and E.S.P. Bromfield 2003). Surprisingly, although we sampled across a wide geographic range, there was no evidence of population structure within each *Ensifer* species. When we assessed isolation by distance separately in *E. medicae* and *E. meliloti,* we failed to detect spatial genetic structure within either rhizobia species in the chromosome or plasmids.

A previous study, which also failed to detect population genetic structure within *Ensifer* species on a large geographic scale, attributed their result to high gene flow among *Ensifer* populations (Silva et al. 2007). High gene flow may explain the lack of genetic structure within *Ensifer* species that we observed as well. The absence of structure across large geographic distances in both studies suggests that dispersal over distances of tens or hundreds of kilometers may frequently occur in *Ensifer.* In addition to this possibility, our data suggest that an absence of genetic structure within *Ensifer* species may be due to limited genetic variation within each species. Nucleotide diversity within each species was at least one order of magnitude lower in its introduced range in North America than in its native range (Epstein et al. 2012). Moreover, we found a near total lack of variation at symbiosis loci within *Ensifer* species, indicating that the absence of genetic structure within each *Ensifer* species does not obscure a significant signal of population differentiation at mutualism-associated loci.

A combination of founder effects, genetic bottlenecks, or recent and limited introduction of bacterial strains likely explains the lack of variation within *Ensifer* species in North America. First, the *Ensifer* samples we collected could be clones of a single strain present in North America. Perhaps a single strain of each *Ensifer* species established in North America when *Ensifer* was introduced in the 1700s (Turkington and Cavers 1979). Alternatively, the strains we sampled could be recent immigrants from *Ensifer’s* native range that have recently displaced older strains. Third, the facultative nature of the *Ensifer-Medicago* interaction may lead to periodic bottlenecks due to strong over-winter selection in the soil that leaves behind limited strains that are capable of associating with plants the following spring. Finally, because we sampled nodules, we only sequenced rhizobium strains that are compatible with *M. lupulina.* Knowing whether the host-compatible rhizobia are only a subset of the diversity of the entire community, as in *Bradyrhizobium* (Sachs et al. 2009), would require a much larger sample of soil diversity. Nevertheless, such a pattern would simply shift the question to why there is such little nucleotide variation among just the compatible strains.

Variation in performance among partner genotypes is important for driving the evolution of partner choice, host sanctions, and cheating in mutualisms, an area that has been explored extensively in the legume-rhizobia symbiosis (Sachs and Simms 2008; Frederickson 2013; Simonsen and Stinchcombe 2014b; Jones et al. 2015). Much of the legume-rhizobia literature assumes that legume plants have a plethora of genetically distinct rhizobia strains to choose from, and that bacterial variation is abundant due to their generation time, numerical abundance, and the number of progeny produced. The relative lack of nucleotide variation within *Ensifer* species — either genome-wide, or in genes implicated in the symbiosis pathway — suggests that in North America the only genetic variation available for plants to select upon is between the two *Ensifer* species. It is possible that recent host-mediated selection reduced diversity within bacteria species, but it is unlikely that such selection would be strong enough to eliminate 99.8% of sequence variation (π values suggests a maximum of 0.1-0.2% sequence variation; Table 1) across a geographic range of ~ 800 km. Nucleotide variation may also be a poor proxy for the quantitative trait variation upon which selection acts. Experimental manipulation of the *Ensifer* symbionts is necessary to explore whether there are differences in the nitrogen fixation efficiency of the two species that might drive local adaptation in the plant host, and evaluate whether genetically divergent *M. lupulina* populations are adapted to different species of rhizobia.

Many classic coevolutionary geographic mosaics comprise only two interacting species (e.g., Brodie et al. 2002). However, geographic mosaics can also involve multispecies assemblages that change in composition across a focal species' range, a pattern documented repeatedly in plant-pollinator mutualisms (e.g., Nagano et al. 2014, Newman et al. 2015). In these systems, spatial variation in pollinator community composition drives corresponding geographic variation in selection on floral phenotypes. The turnover in *Ensifer* assemblages that we observed in the *Medicago-*rhizobia mutualism fits a multispecies view of geographic mosaics. Our data highlight why it is crucial that studies exploring geographic variation in species interactions accurately capture the species assemblages involved. Although most *M. lupulina* plants in Ontario are associated with a different *Ensifer* species than plants in the southeastern United States, we would have concluded that there is no variation in *M. lupulina's* rhizobial partners if we had analyzed each *Ensifer* species independently.

### *Concordant spatial genetic structure in the* M. lupulina *and* Ensifer *mutualism*

A combination of population genetic analyses – isolation by distance, maximum likelihood trees, and population structure analysis – showed strong evidence of genetic differentiation in *M. lupulina* that is somewhat concordant with geographic turnover in *Ensifer* species. We found that *E. meliloti* and *E. medicae* generally occupy different portions of *M. lupulina’s* introduced range. The two *M. lupulina* InStruct clusters weakly correspond to the two *Ensifer* Structure clusters representing the two rhizobia species (Figure 5), and our F_ST_ analysis showed significant genetic differentiation in plants hosting alternative bacterial species. Partially concordant patterns of spatial genetic variation between *Medicago* and the *Ensifer* genus indicate that gene flow could contribute to the maintenance of variation in this mutualism.

In interactions between two partners, gene flow between divergent populations can maintain variation in traits mediating the interaction in both species. In multispecies assemblages—like the *Ensifer* assemblages we documented here—the implications for the maintenance of variation are somewhat different. Gene flow between rhizobia populations is unlikely to introduce new genetic variants within each *Ensifer* species because there is no geographic structure and no genetic variation in either *E. medicae* or *E. meliloti.* Instead, dispersal of *Ensifer* species between populations may maintain variation in rhizobial species diversity in North America. Turnover in *Ensifer* assemblages could contribute to the maintenance of variation in *M. lupulina.* Because *M. lupulina* interacts with two different rhizobia species in eastern North America, gene flow between plant populations partnered with alternate *Ensifer* species has the potential to introduce novel variation in plant mutualism traits. While turnover in *Ensifer* community assemblages may contribute to the maintenance of variation in *M. lupulina* on a continental scale, it is unlikely to be the main source of genetic variation within populations because neighboring populations tend to have the same species of *Ensifer.*

There is suggestive evidence that genetic differentiation among *Medicago* populations may arise in part from geographically structured coevolution with *Ensifer* assemblages. Béna et al. (2005)found evidence that geographically structured diversity in rhizobia potentially influenced geographically structured diversity in the *Medicago* genus in its native Eurasian range. Population genetic differentiation in *Medicago* could result from adaptation to local strains that differ in nitrogen fixation effectiveness. The *E. medicae* lab strain WSM419 is a more effective mutualist than the lab strain *E. meliloti* 1021 (Terpolilli et al. 2008), which if generally true of these species, suggests that the *Ensifer* species common in southern Ontario populations is more effective than the *Ensifer* species common in the southeastern United States. However, it is not necessarily appropriate to extrapolate these lab results to genetically heterogeneous natural populations, given that Béna et al. (2005) showed that rhizobia effectiveness is contingent on the specific legume host, and Terpolilli et al. (2008) evaluated the *Ensifer* species with a single *M. truncatula* genotype.

Concordant genetic structure in interacting species may not arise from coevolutionary processes that maintain genetic variation and facilitate future coevolution. The genetic differences between *M. lupulina* populations and geographic turnover in *Ensifer* assemblages could be due to several other processes, including multiple introductions to North America, adaptation to other aspects of the environment, or neutral forces. Local adaptation to the substantial climatic differences between southern Ontario and the southeastern United States (e.g., temperature, precipitation) could contribute to geographic structure in both *Medicago* and *Ensifer.* In addition, *Ensifer* associations with other *Medicago* species in North America, such as *M. sativa* and *M. polymorpha* (Béna et al. 2005; Rome et al. 1996), could be driving large-scale patterns in *Ensifer* species distribution. Genetic structure in *M. lupulina* in its native range has been attributed to self-fertilization (Yan et al. 2009), and likely contributes to the isolation by distance we observed as well. Evaluating the mechanisms behind the geographic trends that we observed is a separate question from the maintenance of genetic variation that ultimately requires manipulative field experiments that are logistically challenging to perform with bacteria (but see Simonsen and Stinchcombe, 2014a). Despite these alternative explanations for the somewhat concordant patterns of geographic structure in *M. lupulina* and its rhizobial mutualist *Ensifer*, the significant potential for coevolution between *M. lupulina* and *Ensifer* assemblages we discovered in this study is worth further investigation. Future work involving experiments testing local adaptation of *M. lupulina* plants to its local *Ensifer* species could reveal additional evidence of coevolution in this system in the its introduced range in North America.

### Conclusions and Prospects

Comparing spatial genetic structure and genome-wide variation in mutualist partners is an effective approach to determine the potential scale of coevolution between interacting species. Given the relative lack of genome-wide variation within *E. medicae* and *E. meliloti,* differences between *Ensifer* species are the only potential source of coevolutionary selection acting on *M. lupulina.* Our study shows how comparing geographic variation in two mutualists is important to understand how variation may be maintained in mutualisms, especially in introduced ranges where processes like gene flow, bottlenecks, and multiple introductions can complicate mutualist interactions.

## Acknowledgements

We thank Stephen Wright, Nicole Mideo, Benjamin Gilbert, and Megan Frederickson for comments and discussion, Andrew Hall and Bruce Petrie for plant growth assistance, and Maggie Bartkowsksa, Adriana Salcedo, and Billie Gould for bioinformatics advice. Our work is supported by NSERC Discovery Grants and Graduate Scholarships (JRS, TLH), an EEB Departmental Post-Doc Fellowship (CWW), and the National Science Foundation (KDH). We are grateful to Mohamed Noor, Maurine Neiman, and two anonymous reviewers for comments that greatly improved this manuscript.

## Data accessibility

Sequence data will be made available on NCBI. Input VCF files, Python script, and R scripts will be made available on Dryad (DOI number will be finalized upon acceptance). Sampling locations are available in Table 1 of the Supplemental Materials.

**References**

